# Cyclophilin A Prevents HIV-1 Restriction in Lymphocytes by Blocking Human TRIM5α Binding to the viral Core

**DOI:** 10.1101/678037

**Authors:** Anastasia Selyutina, Mirjana Persaud, Angel Bulnes-Ramos, Cindy Buffone, Alicia Martinez-Lopez, Viviana Scoca, Francesca Di Nunzio, Joseph Hiatt, Nevan J. Krogan, Judd F. Hultquist, Felipe Diaz-Griffero

## Abstract

Disruption of cyclophilin A (CypA)-capsid interactions affects HIV-1 replication in human lymphocytes. To understand the mechanism, we used Jurkat cells, human PBMCs, and human CD4^+^ T cells. Our results showed that the inhibition of HIV-1 infection caused by disrupting CypA-capsid interactions is dependent on human TRIM5α (TRIM5α_hu_), suggesting that TRIM5α_hu_ restricts HIV-1. Accordingly, we found that TRIM5α_hu_ binds to the HIV-1 core. Disruption of CypA-capsid interactions failed to affect HIV-1-A92E infection, correlating with the loss of TRIM5α_hu_ binding to HIV-1-A92E cores. Disruption of CypA-capsid interactions in PBMCs and CD4^+^ T cells had a greater inhibitory effect on HIV-1 when compared to Jurkat cells. HIV-1-A92E infection of PBMCs and CD4^+^ T cells was unaffected by disruption of CypA-capsid interactions. Consistent with TRIM5α restriction, disruption of CypA-capsid interactions in CD4^+^ T cells inhibited reverse transcription. Overall, our results showed that CypA binding to the core protects HIV-1 from TRIM5α_hu_ restriction.

## INTRODUCTION

The early steps of human immunodeficiency virus (HIV-1) replication involve delivery of the viral core into the host cytoplasm. The viral core is a conical structure composed of ∼1,500 monomers of the HIV-1 capsid protein, which protects the viral RNA genome (Pornillos et al., 2009; Sundquist and Hill, 2007). The capsid protein plays a role in uncoating, reverse transcription, and nuclear import. To perform these functions, the capsid protein has been shown to functionally interact with several host dependency factors, including cyclophilin A (CypA) (Luban et al., 1993), Nup153 (Buffone et al., 2018; Di Nunzio et al., 2013) (Brass et al., 2008; Konig et al., 2008; Lee et al., 2010; Matreyek and Engelman, 2011; Matreyek et al., 2013; Zhou et al., 2008), RANBP2 (Di Nunzio et al., 2012; Meehan et al., 2014; Ocwieja et al., 2011), and TNPO3 (Brass et al., 2008; Christ et al., 2008; Konig et al., 2008; Krishnan et al., 2010; Levin et al., 2010; Ocwieja et al., 2011; Thys et al., 2011; Valle-Casuso et al., 2012; Zhou et al., 2008). Effective HIV-1 infection may occur only when these factors interact with the HIV-1 core at the appropriate time and in the appropriate order, thus raising the possibility that the initial binding of particular factors may modulate the subsequent binding of other factors.

The interaction of CypA with monomeric HIV-1 capsid was originally reported about 30 years ago (Luban et al., 1993), and it has been well-established that this interaction modulates HIV-1 infection (Braaten et al., 1996; Braaten and Luban, 2001; Li et al., 2009; Schaller et al., 2011; Sherry et al., 1998; Sokolskaja et al., 2004; Thali et al., 1994; Towers et al., 2003). Although the ability of CypA to modulate HIV-1 infection is cell-type dependent, it is clear that the disruption of CypA-capsid interactions negatively affects HIV-1 infection of lymphocytes (Braaten and Luban, 2001; Li et al., 2009; Ptak et al., 2008; Sherry et al., 1998; Sokolskaja et al., 2004; Towers et al., 2003; Wainberg et al., 1988; Yin et al., 1998); however, the underlying mechanism remains elusive.

To investigate the molecular events that result in a decrease in HIV-1 infection upon disruption of CypA-capsid interactions in lymphocytes, we used Jurkat cells, primary human peripheral blood mononuclear cells (PBMCs), and primary human CD4^+^ T cells as experimental models. Our experiments revealed that inhibition of HIV-1 infection caused by disrupting CypA-capsid interactions is dependent on the expression TRIM5α_hu_. These results suggested that TRIM5α_hu_ is restricting HIV-1 infection upon disruption of CypA-capsid interactions. In agreement, we found that TRIM5α_hu_ binds to the HIV-1 core. Interestingly, TRIM5α_hu_ does not bind to HIV-1 cores bearing the capsid mutant A92E (HIV-1-A92E). Infection of HIV-1-A92E viruses was not affected by disruption of CypA-capsid interactions, which correlates with the loss of TRIM5α_hu_ binding to HIV-1 cores bearing the A92E mutation. These experiments suggested that CypA protects the HIV-1 core from TRIM5α_hu_. Although the infectivity defect observed in Jurkat cells is small, disruption of CypA-interactions in PBMCs and CD4^+^ T cells causes a major defect in infectivity. In agreement with our hypothesis that CypA is protecting the core from TRIM5α_hu_, infection of PBMCs and CD4^+^ T cells by HIV-1-A92E viruses was insensitive to the disruption of CypA-capsid interactions. Consistently with a block imposed by TRIM5α proteins, disruption of CypA-capsid interactions in CD4^+^ T cells dramatically decreased the efficiency of reverse transcription. Overall, our investigations suggested a model in which CypA binding to the HIV-1 core protects the core from the restriction factor TRIM5α_hu_.

## RESULTS

### HIV-1 infection of Jurkat cells is sensitive to Cyclosporin A treatment

To understand the role of CypA in HIV-1 infection of CD4^+^ T cells, we used the human T-cell line Jurkat as an experimental model. Cyclosporin A (CsA) is a molecule that binds to CypA, and thus blocks cellular CypA activity. To measure the effect of CsA treatment on HIV-1 infectivity, we infected Jurkat cells with HIV-1 tagged with green fluorescent protein (HIV-1-GFP) in the presence of 10 µM CsA. Consistent with results from previous studies (Braaten and Luban, 2001; Hatziioannou et al., 2005; Li et al., 2009; Ptak et al., 2008; Sokolskaja et al., 2004; Towers et al., 2003; Wainberg et al., 1988; Yin et al., 1998), HIV-1-GFP infection of human Jurkat T cells was sensitive to CsA treatment (Figure 1A, upper panel). Experiments were repeated multiple times and the fraction of GFP-positive cells normalized to the dimethyl sulfoxide (DMSO)-treated control is shown (Figure 1A, lower panel). To further confirm the importance of CypA-capsid interactions in HIV-1 infection, we challenged Jurkat T cells with p24-normalized amounts of HIV-1-P90A and HIV-1-G89V, which are mutant versions of HIV-1 that disrupt CypA-capsid interactions (Gamble et al., 1996; Thali et al., 1994). Infection was determined by measuring the percentage of GFP-positive cells at 48 h post-infection. We found that the mutant HIV-1 strains bearing capsid changes that disrupted CypA-capsid interactions were less infectious than wild-type HIV-1 (Figure 1B, upper panel). Experiments were repeated multiple times and the fraction of GFP-positive cells normalized to control is shown (Figure 1B, lower panel). Although Jurkat cells failed to show a dramatic decrease in HIV-1 infectivity as a result of disrupted CypA-capsid interactions, we still used this cell line as a model to mechanistically understand the contribution of CypA to HIV-1 infection.

**Figure 1.**
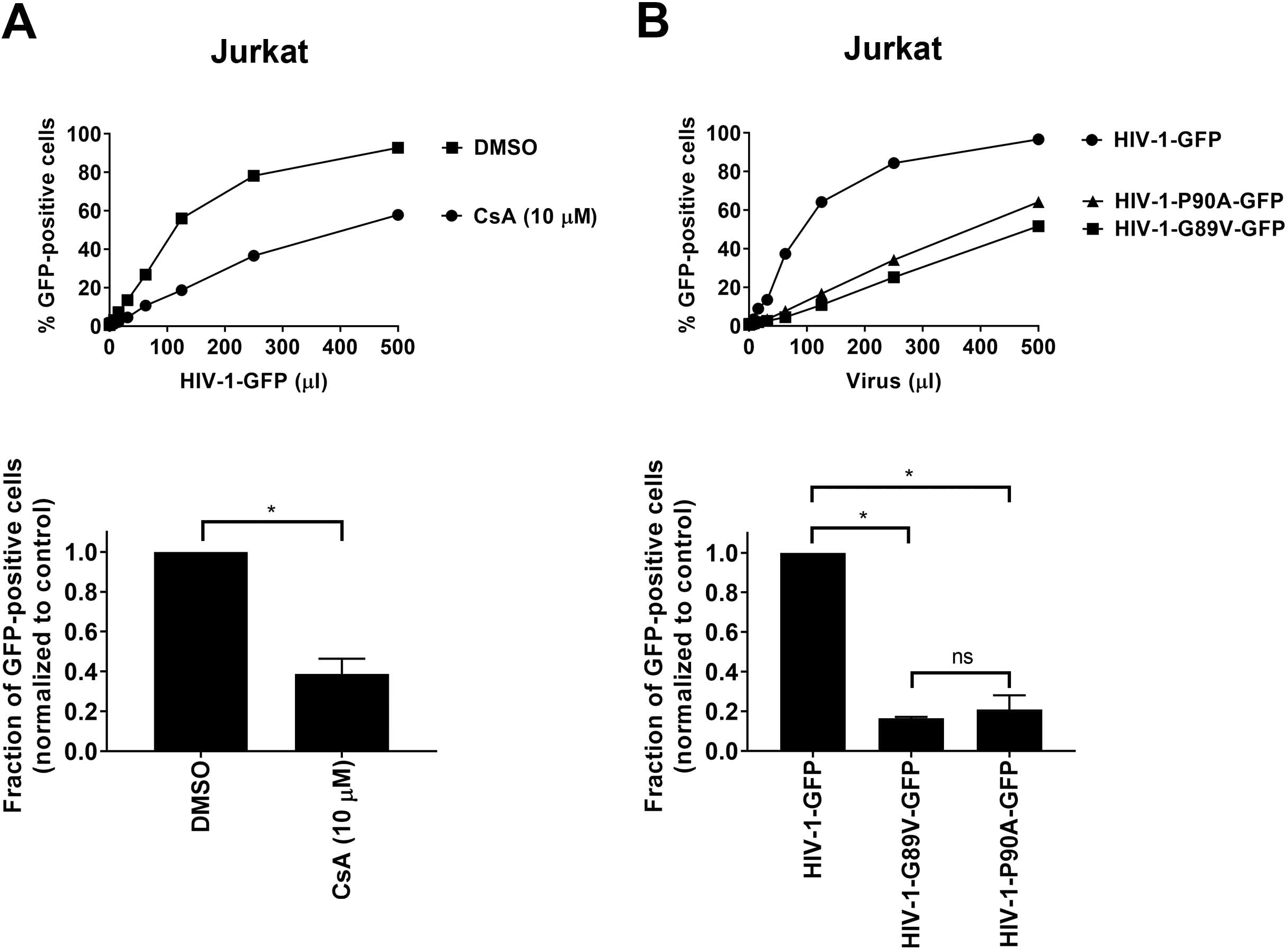
CypA is important for HIV-1 infection of Jurkat cells. **(A)** Human Jurkat cells were challenged with increasing amounts of HIV-1 expressing GFP as a reporter of infection (HIV-1-GFP) in the presence of 10 µM cyclosporin A (CsA) (upper panel). DMSO was used as control. Infection was determined at 48 h post-challenge by measuring the percentage of GFP-positive cells using a flow cytometer. Similar results were obtained from three independent experiments and a representative experiment is shown. The result of three independent experiments normalized to wild-type HIV-1-GFP infection with standard deviation is shown (lower panel). * indicates P-value < 0.005 as determined by the unpaired t-test. **(B)** Jurkat cells were challenged with increasing amounts of p24-normalized HIV-1, HIV-1-P90A, or HIV-1-G89V expressing GFP (upper panel). Infection was determined at 48 h post-challenge by measuring the percentage of GFP-positive cells using a flow cytometer. Similar results were obtained from three independent experiments and a representative experiment is shown. The results of three independent experiments normalized to wild-type HIV-1-GFP infection with standard deviation is shown (lower panel). * indicates P-value < 0.005, ns indicates not significant as determined by the Two-way ANOVA Tukey’s multiple comparisons test.

### Human TRIM5α (TRIM5α_hu_) binds to the HIV-1 capsid

Previous studies have shown that CypA modulates the ability of rhesus tripartite motif (TRIM) protein, TRIM5α (TRIM5α_rh_), to restrict HIV-1 infection (Berthoux et al., 2005; Burse et al., 2017; Keckesova et al., 2006; Lin and Emerman, 2008; Stremlau et al., 2006b), which is not surprising because both proteins interact with the HIV-1 core. It is well known that TRIM5α_rh_ cages the HIV-1 core using cooperative binding during infection (Diaz-Griffero et al., 2009; Ganser-Pornillos et al., 2011; Stremlau et al., 2006a). Because both CypA and TRIM5α_rh_ bind to the HIV-1 core, it is possible that CypA modulates the binding of TRIM5α_rh_ to the HIV-1 core and vice versa.

More recent studies have suggested that TRIM5α_hu_ blocks HIV-1 infection (Jimenez-Guardeno et al., 2019; OhAinle et al., 2018), possibly by binding to the HIV-1 core. Therefore, we tested the ability of TRIM5α_hu_ to bind to the HIV-1 core using our capsid binding assay (Selyutina et al., 2018). We found that TRIM5α_hu_ binds to the HIV-1 core (Figure 2). TRIM5α_hu_ showed a stronger interaction with the HIV-1 core compared with TRIM5α_rh_ (Figure 2). Our results showed for the first time that TRIM5α_hu_ interacted with the HIV-1 core. One possibility is that CypA modulates the ability of TRIM5α_hu_ to interact with the HIV-1 core.

**Figure 2.**
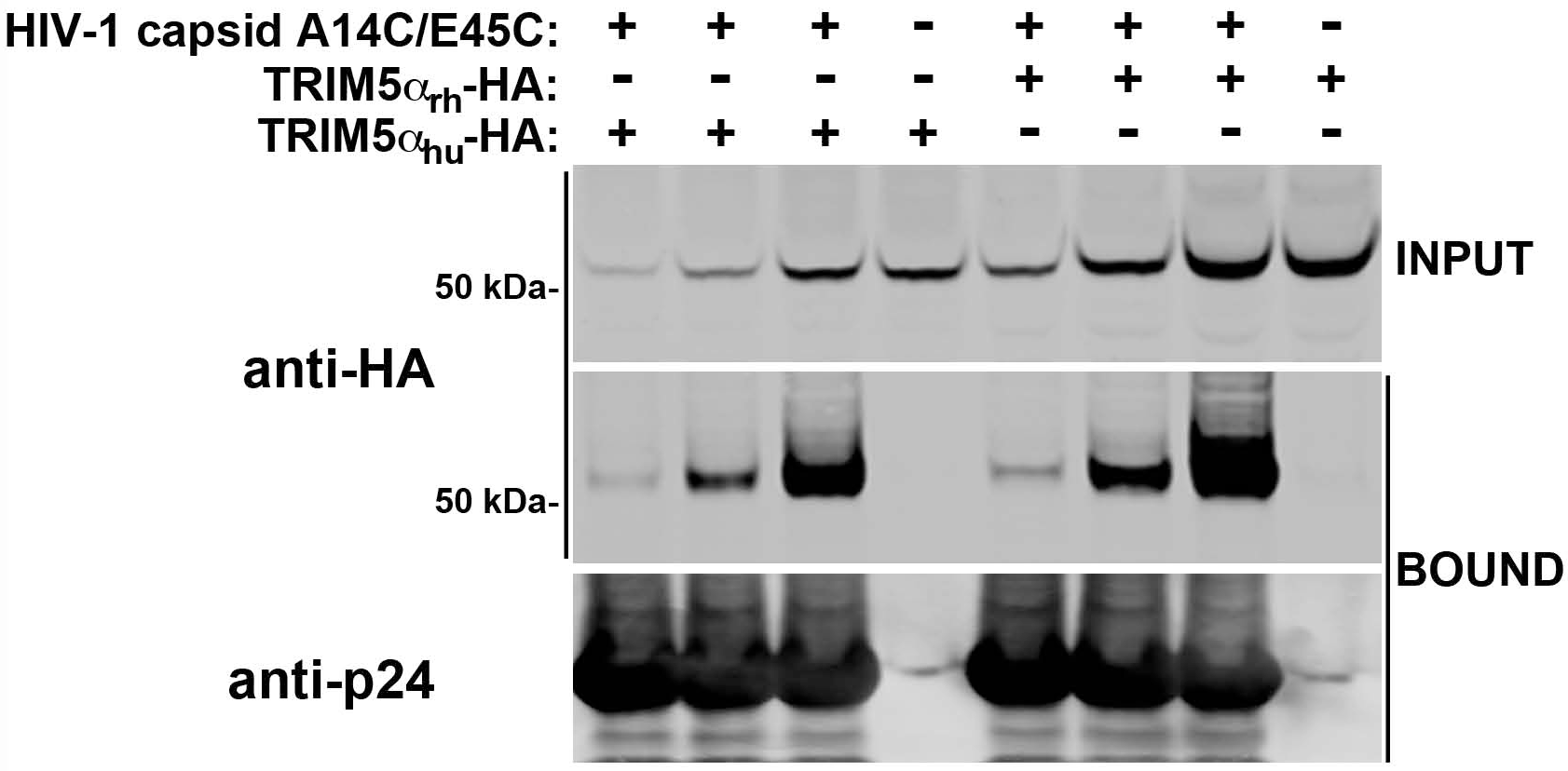
TRIM5α_hu_ binds to the HIV-1 core. Human 293T cells were transfected with plasmids expressing the wild-type human TRIM5α tagged with an HA epitope (TRIM5α_hu_-HA). At 24 h post-transfection, cells were lysed in capsid binding buffer (CBB), as described in Materials and Methods. Inputs containing different amounts of TRIM5α_hu_were prepared (INPUT). Cell extracts containing different amounts of TRIM5α_hu_-HA were mixed with 20 μl of stabilized HIV-1 capsid tubes (5 mg/ml) and incubated for 1 h at room temperature. Stabilized HIV-1 capsid tubes were collected by centrifugation and washed twice using CBB. Pellets were resuspended in 1× Laemmli buffer (BOUND). INPUT and BOUND fractions were analyzed by western blotting using anti-HA and anti-p24 antibodies. As control, similar experiments were performed using the capsid binder, rhesus TRIM5α tagged with an HA epitope (TRIM5α_rh_-HA). Each experiment was repeated at least six times and a representative experiment is shown.

### TRIM5α_hu_ expression is necessary for the inhibition of HIV-1 infection mediated by the disruption of CypA-capsid interactions

Because of the ability of TRIM5α_hu_ to bind to the HIV-1 core, we tested whether TRIM5α_hu_ is involved in the blocking of HIV-1 infection mediated by the disruption of CypA-capsid interactions. For this purpose, we achieved knockdown (KD) of TRIM5α_hu_ expression in Jurkat cells using a specific short hairpin RNA (shRNA) against TRIM5α_hu_ (Figure 3), as described in Materials and Methods. To test for TRIM5α_hu_ expression, we challenged different sets of TRIM5α_hu_ KD cells with increasing amounts of N-tropic murine leukemia virus GFP (N-MLV-GFP), which is restricted by TRIM5α_hu_. We found that Jurkat TRIM5α_hu_ KD cells were susceptible to N-MLV infection (Figure 3A), whereas in shRNA control cells, N-MLV-GFP infection was blocked. As a control, we also showed that TRIM5α_hu_ KD cells were susceptible to B-tropic murine leukemia virus GFP (B-MLV-GFP), which is not restricted by TRIM5α_hu_ (Figure 3A). These results confirmed that KD of TRIM5α_hu_ expression was successfully achieved.

**Figure 3.**
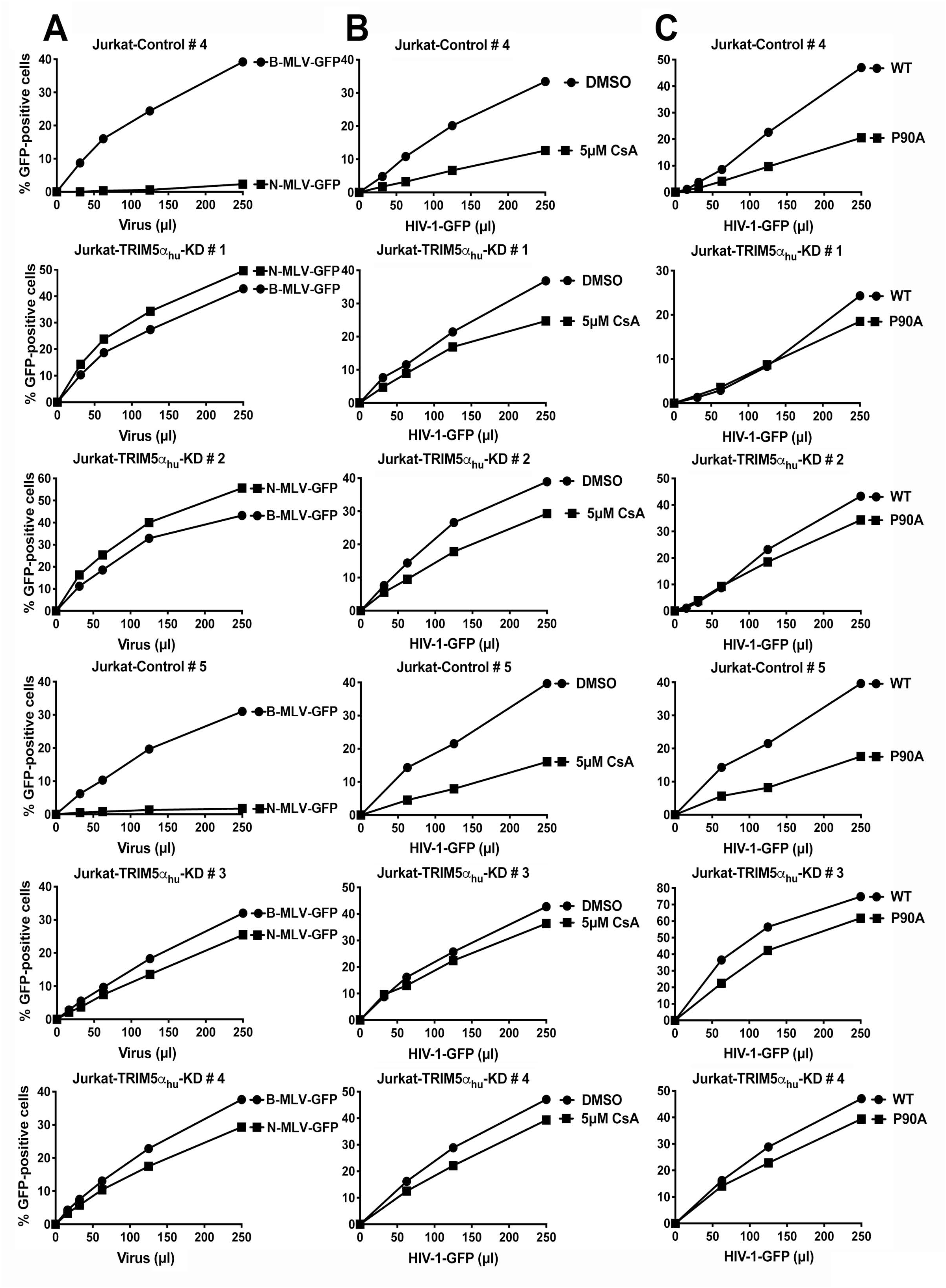
Expression of TRIM5α_hu_ is necessary for the inhibition of HIV-1 infection caused by the disruption of CypA-capsid interactions. **(A)** Human Jurkat cells stably expressing a specific shRNA against TRIM5α_hu_ (Jurkat-TRIM5α_hu_-KD #1–4) or the empty vector (Jurkat-Control #4 and 5) were challenged using increasing amounts of N-MLV-GFP, which are restricted by TRIM5α_hu_. Infection was determined at 48 h post-challenge by measuring the percentage of GFP-positive cells using a flow cytometer. As a control, similar infections were performed using B-MLV-GFP viruses, which are not restricted by TRIM5α_hu_. **(B)** Jurkat-TRIM5α_hu_-knockdown (KD) and Jurkat-control cells were challenged with increasing amounts of HIV-1-GFP in the presence of cyclosporin A (CsA). Infection was determined at 48 h post-challenge by measuring the percentage of GFP-positive cells using a flow cytometer. As a control, similar experiments were performed using DMSO. **(C)** Jurkat-TRIM5α_hu_-KD and Jurkat-control cells were challenged with increasing amounts of p24-normalized HIV-1-GFP or HIV-1-P90A-GFP. Infection was determined at 48 h post-challenge by measuring the percentage of GFP-positive cells using a flow cytometer. Each experiment was repeated at least three times and a representative experiment is shown.

Next, we assessed the infection phenotype observed in Jurkat TRIM5α_hu_ KD cells when CypA-capsid interactions are disrupted (Figure 3B). Surprisingly, TRIM5α_hu_ KD cells treated with CsA did not show marked inhibition of HIV-1 infection compared with control cells (Figure 3B), suggesting that TRIM5α_hu_ is involved in the inhibition of viral infection observed in CsA-treated cells that have disrupted CypA-capsid interactions.

HIV-1-P90A is a mutant form of HIV-1 in which CypA-capsid interactions are disrupted during infection. We tested whether HIV-1-P90A infection was inhibited in TRIM5α_hu_ KD Jurkat cells. As shown in Figure 3C, we found that HIV-1-P90A infection was not inhibited in TRIM5α_hu_ KD cells compared with control cells. This result supports our hypothesis that expression of TRIM5α_hu_ is important for the infection phenotype observed when CypA-capsid interactions are disrupted. It is possible that CypA prevents the binding of TRIM5α_hu_ to the HIV-1 core via steric hindrance, thus protecting the core from the restriction factor during the early steps of infection.

### CypA prevents the interaction of TRIM5α_hu_ with the HIV-1 core

To test whether CypA modulates the binding of TRIM5α_hu_ to the HIV-1 core, we analyzed the binding of TRIM5α_hu_ to the HIV-1 core in the presence of CsA, which binds to endogenously expressed CypA in cellular extracts and prevents CypA binding to the HIV-1 core. As shown in Figure 4A, TRIM5α_hu_ binding to the HIV-1 core increased 3- to 4-fold in the presence of CsA. We also found that the binding of TRIM5α_hu_ to mutant P90A HIV-1 cores was 5- to 6-fold higher than that of wild-type HIV-1 cores (Figure 4A) in the absence or presence of CsA. These results suggested that CypA prevents the binding of TRIM5α_hu_ to the HIV-1 core during infection. It is possible that CypA and TRIM5α_hu_ compete with each other to interact with the incoming core. To test this hypothesis, we investigated whether the addition of exogenous CypA decreased TRIM5α_hu_ binding to the HIV-1 core. For this purpose, we assessed the ability of TRIM5α_hu_ from cellular extracts to bind to the HIV-1 core in the presence of 10 µg of recombinant CypA protein. We found that the addition of recombinant CypA significantly decreased the binding of TRIM5α_hu_ to the HIV-1 core (Figure 4B). As expected, recombinant CypA did not affect the binding of TRIM5α_hu_ to HIV-1 cores bearing the P90A mutation. Overall, these results suggested that CypA protects the core from the restriction factor TRIM5α_hu_.

**Figure 4.**
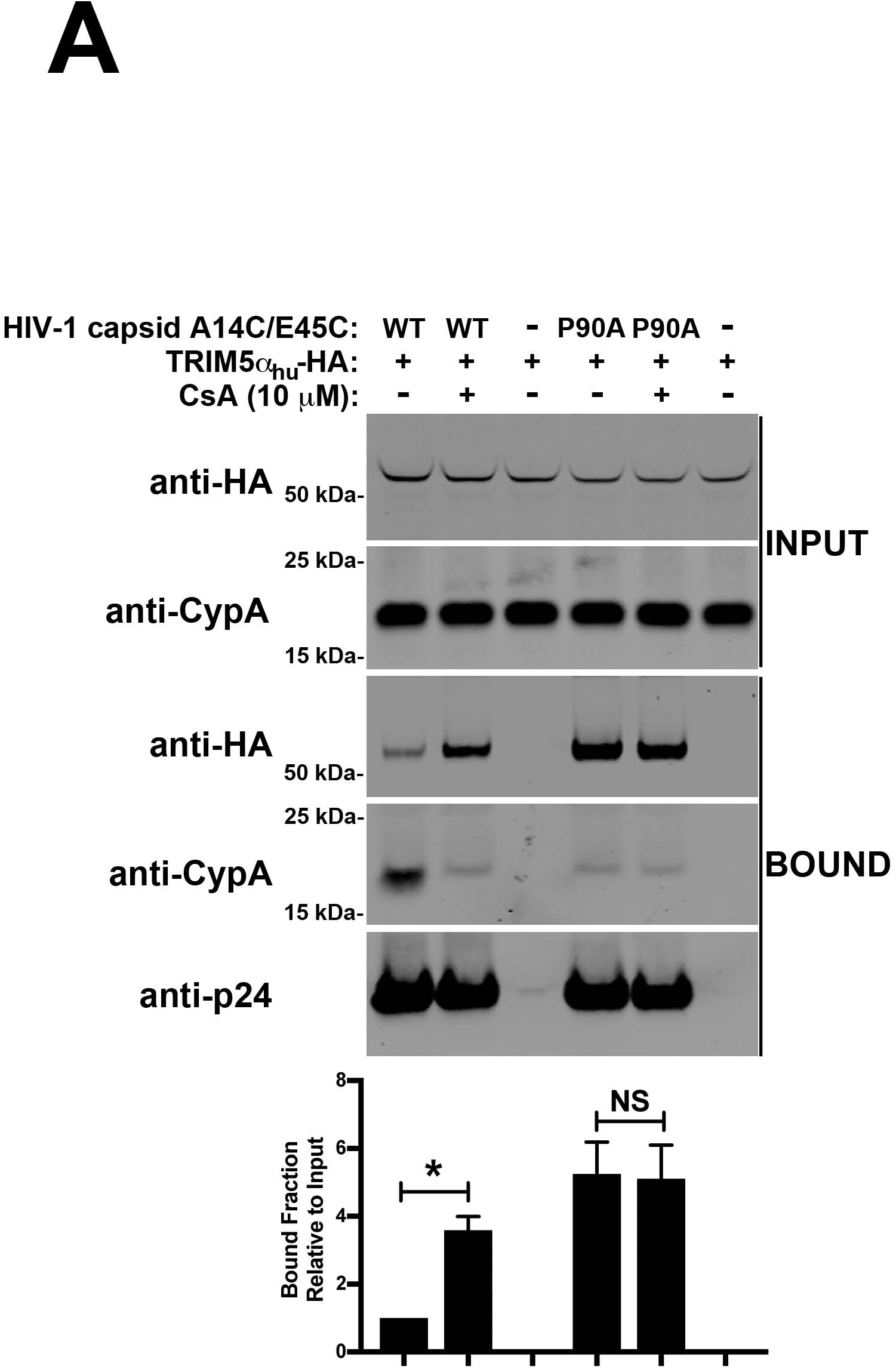
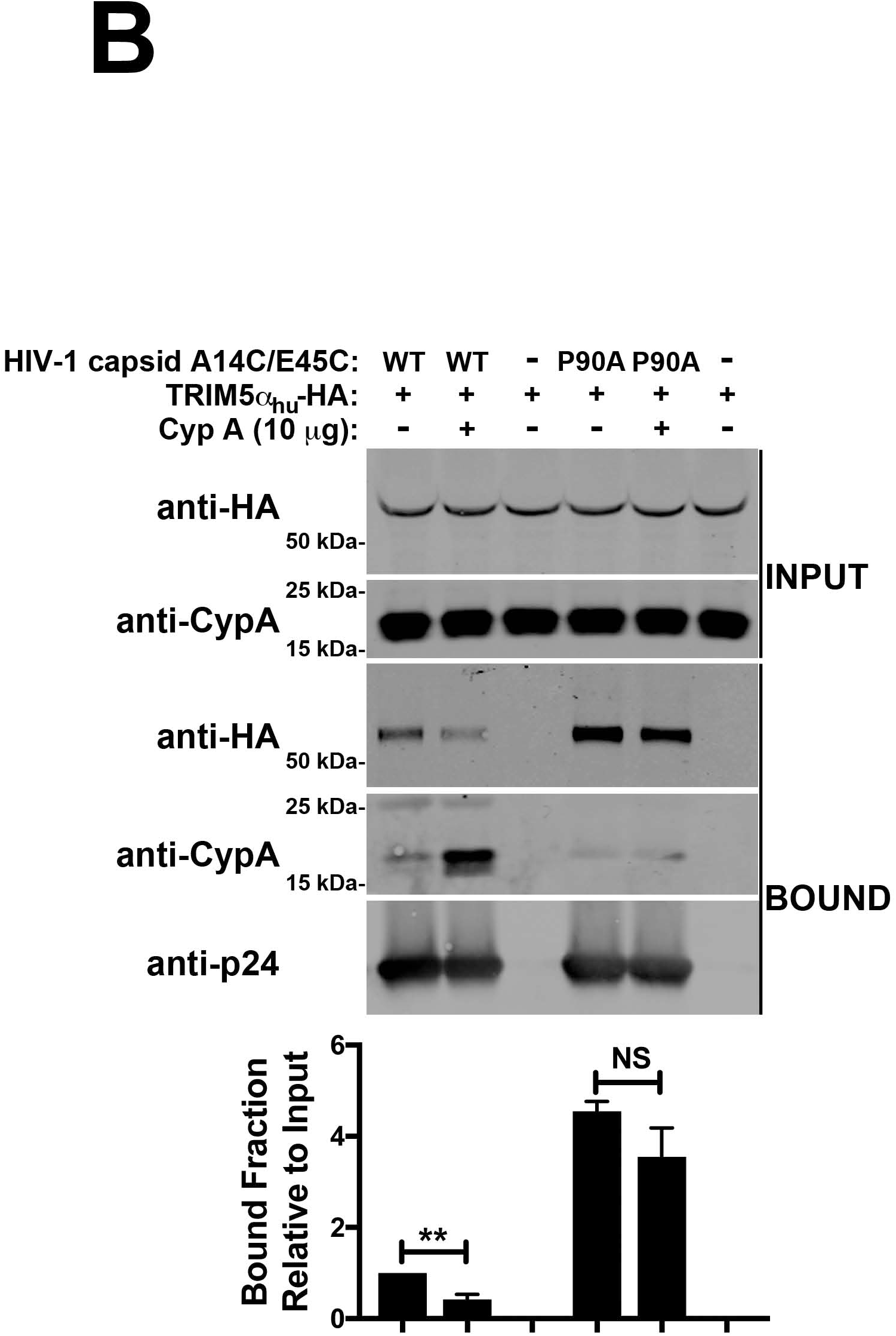
CypA prevents the binding of TRIM5α_hu_ to the HIV-1 core. **(A)** Human 293T cells were transfected with plasmids expressing TRIM5α_hu_-HA. At 24 h post-transfection, cells were lysed in capsid binding buffer (CBB) (INPUT). Cell extracts containing TRIM5α_hu_-HA were mixed with 20 μl of stabilized wild-type or P90A HIV-1 capsid tubes (5 mg/ml) in the presence of 10 µM CsA. Mixtures were incubated for 1 h at room temperature. Stabilized HIV-1 capsid tubes were collected by centrifugation and washed twice using CBB. Pellets were resuspended in 1X Laemmli buffer (BOUND). INPUT and BOUND fractions were analyzed by western blotting using anti-HA, anti-CypA, and anti-p24 antibodies. Experiments were repeated three times and a representative experiment is shown. The bound fraction relative to input for three independent experiments with standard deviation is shown. * indicates P-value < 0.005, ns indicates not significant as determined by the Two-way ANOVA Tukey’s multiple comparisons test. **(B)** Cell extracts containing TRIM5α_hu_-HA were mixed with 20 μl of stabilized wild-type or P90A HIV-1 capsid tubes (5 mg/ml) in the presence of 10 µg recombinant CypA. Mixtures were incubated for 1 h at room temperature. Stabilized HIV-1 capsid tubes were collected by centrifugation and washed twice using CBB. Pellets were resuspended in 1× Laemmli buffer (BOUND). INPUT and BOUND fractions were analyzed by western blotting using anti-HA, anti-CypA, and anti-p24 antibodies. Experiments were repeated three times and a representative experiment is shown. The bound fraction relative to input for three independent experiments with standard deviation is shown. * indicates P-value < 0.005, ns indicates not significant as determined by the Two-way ANOVA Tukey’s multiple comparisons test.

### Mutant HIV-1 strains that are impaired for capsid-TRIM5α_hu_ interactions are insensitive to CsA treatment

Previous studies have shown that infection of Jurkat cells by the HIV-1 strain bearing the capsid mutation A92E (HIV-1-A92E-GFP) was less sensitive to CsA treatment when compared with wild-type HIV-1 (Hatziioannou et al., 2005; Sokolskaja et al., 2004). Consistent with these findings, we observed that HIV-1-A92E infection of Jurkat cells was insensitive to CsA treatment (Figure 5A). This insensitivity may occur due to one or both of the following scenarios: 1) HIV-1-A92E may have a higher affinity for CypA, which may better protect the virus from TRIM5α_hu_, and 2) HIV-1-A92E may have a lower affinity for TRIM5α_hu_, which may prevent inhibition of infection. To explore these possibilities, we tested the ability of CypA to interact with HIV-1 cores containing the A92E capsid mutation. We found that CypA did not bind to the mutant A92E core, which suggested no or poor interaction between HIV-1-A92E and CypA during infection (Figure 5B). As a control, we showed that the binding of CypA to wild-type HIV-1 cores was CsA-sensitive (Figure 5B). Because CypA does not bind to HIV-1-A92E, a stronger binding between HIV-1-A92E and TRIM5α_hu_ may result in inhibition of infection. However, we found that the infection of Jurkat cells by HIV-1-A92E was not affected when compared with that of wild-type HIV-1 (Figure 5A), suggesting that HIV-1-A92E may be unable to interact with TRIM5α_hu_. Consistent with our hypothesis, TRIM5α_hu_ did not interact with mutant HIV-1 cores bearing the capsid change A92E (Figure 5C). Overall, these results indicated that mutant HIV-1 A92E cores bind neither CypA nor TRIM5α_hu,_ which may explain the reason that CsA treatment does not affect HIV-1-A92E infection of Jurkat cells.

**Figure 5.**
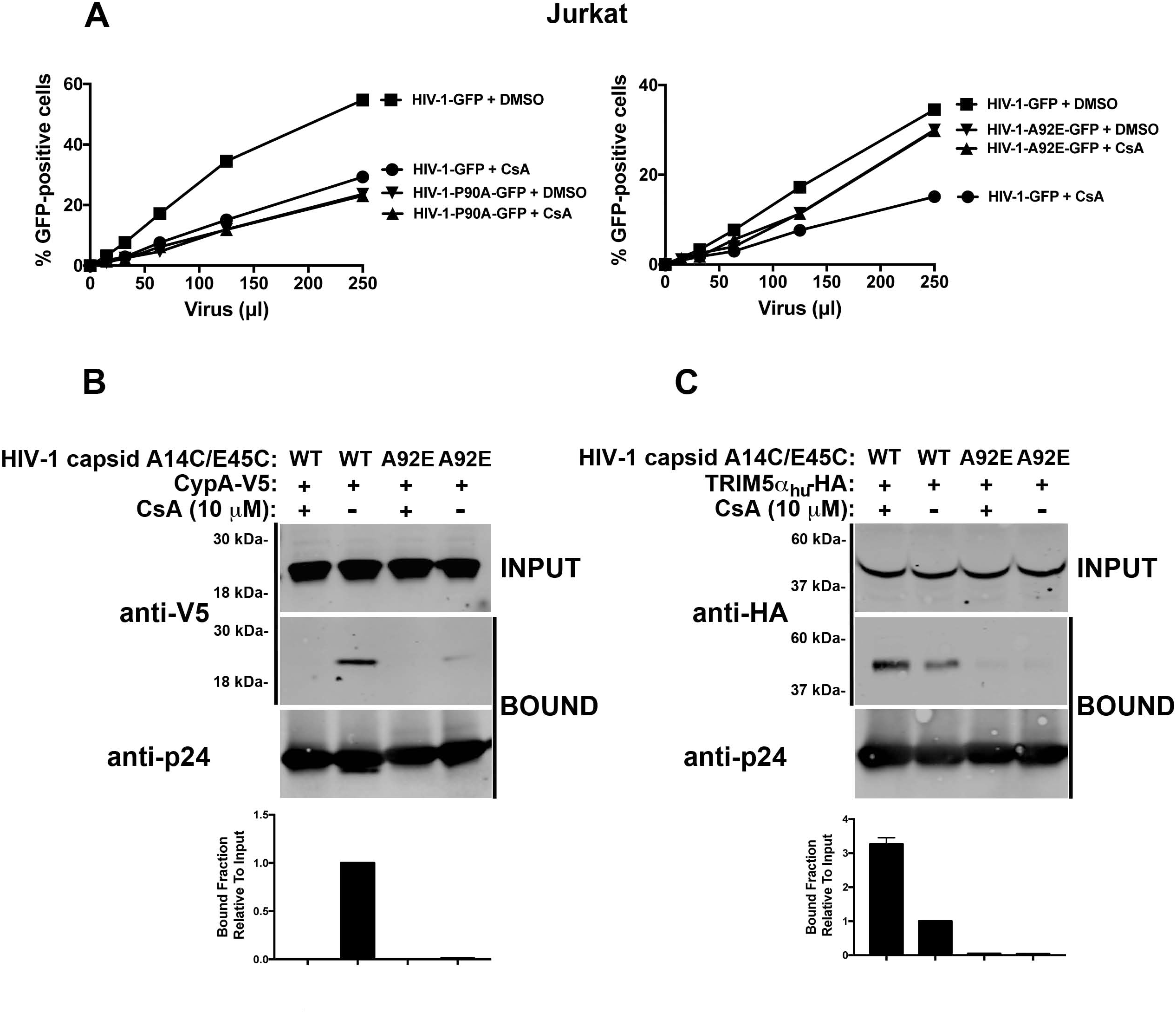
HIV-1 viruses bearing a capsid mutant that does not interact with TRIM5α_hu_ are insensitive to CsA. **(A)** HIV-1-A92E infection of Jurkat cells. Jurkat cells were challenged with increasing amounts of p24-normalized HIV-1-GFP, HIV-1-P90A-GFP (left panel), or HIV-1-A92E-GFP (right panel) in the presence of 10 μM Cyclosporin A (CsA). DMSO was used as a negative control. Infection was determined at 48 h post-challenge by measuring the percentage of GFP-positive cells using a flow cytometer. Experiments were repeated three times and a representative experiment is shown. **(B)** Binding of CypA to HIV-1 cores bearing the capsid change A92E. Cell extracts from human 293T cells containing CypA-V5 were mixed with 20 μl of stabilized wild-type or A92E HIV-1 capsid tubes (5 mg/ml) in the presence of 10 µM CsA. Mixtures were incubated for 1 h at room temperature. Stabilized HIV-1 capsid tubes were collected by centrifugation and washed twice using capsid binding buffer (CBB). Pellets were resuspended in 1× Laemmli buffer (BOUND). INPUT and BOUND fractions were analyzed by western blotting using anti-V5 and anti-p24 antibodies. Experiments were repeated three times and a representative experiment is shown. The bound fraction relative to input for three independent experiments with standard deviation is shown. **(C)** Binding of TRIM5α_hu_ to HIV-1 cores bearing the capsid change A92E. Similarly, cell extracts containing TRIM5α_hu_-HA were mixed with 20 μl of stabilized wild-type or A92E HIV-1 capsid tubes (5 mg/ml) in the presence of 10 µM CsA. Mixtures were incubated for 1 h at room temperature. Stabilized HIV-1 capsid tubes were collected by centrifugation and washed twice using CBB. Pellets were resuspended in 1× Laemmli buffer (BOUND). INPUT and BOUND fractions were analyzed by western blot using anti-HA and anti-p24 antibodies. Experiments were repeated three times and a representative experiment is shown. The bound fraction relative to input for three independent experiments with standard deviation is shown.

### CypA-capsid interactions are required for HIV-1 infection of human PBMCs and CD4^+^ T cells

To understand the role of CypA in HIV-1 infection of human primary PBMCs, we challenged PBMCs and CD4^+^ T cells from different donors with HIV-1 in the presence of increasing concentrations of CsA, which prevents the binding of CypA to HIV-1 capsid (Diaz-Griffero et al., 2006; Luban et al., 1993). As shown in Figure S1, CsA treatment inhibited HIV-1 infection in PBMCs and CD4^+^ T cells. This result suggested that CypA-capsid interactions are required for HIV-1 infection of human PBMCs and CD4^+^ T cells. To further corroborate these findings, we challenged human PBMCs and CD4^+^ T cells with increasing amounts of HIV-1 particles bearing the capsid mutations P90A and G89V, both of which prevent CypA-capsid interactions (Gamble et al., 1996; Thali et al., 1994). Infection was determined by measuring the percentage of GFP-positive cells at 48 h post-infection. Wild-type and mutant HIV-1-GFP viruses were normalized by quantifying the levels of p24. We found that HIV-1-P90A and HIV-1-G89V were unable to infect human PBMCs and CD4^+^ T cells (Figure 6A and B). Although this experiment was performed in PBMCs and CD4^+^ T cells obtained from six different donors, all of which showed the same results, we have only shown sample results obtained from three donors. Our results suggested that CypA-capsid interactions are required for HIV-1 infection in human primary PBMCs and CD4^+^ T cells. Importantly, the lack of CypA appeared to have a stronger effect on HIV-1 infection in primary human lymphocytes compared with that in Jurkat cells (Figures 1B and 6A&B), thus indicating that CypA was crucial for HIV-1 infection of human primary cells. Overall, these results suggested that the disruption of CypA-capsid interaction is detrimental for HIV-1 infection of PBMCs and CD4^+^ T cells.

**Figure 6.**
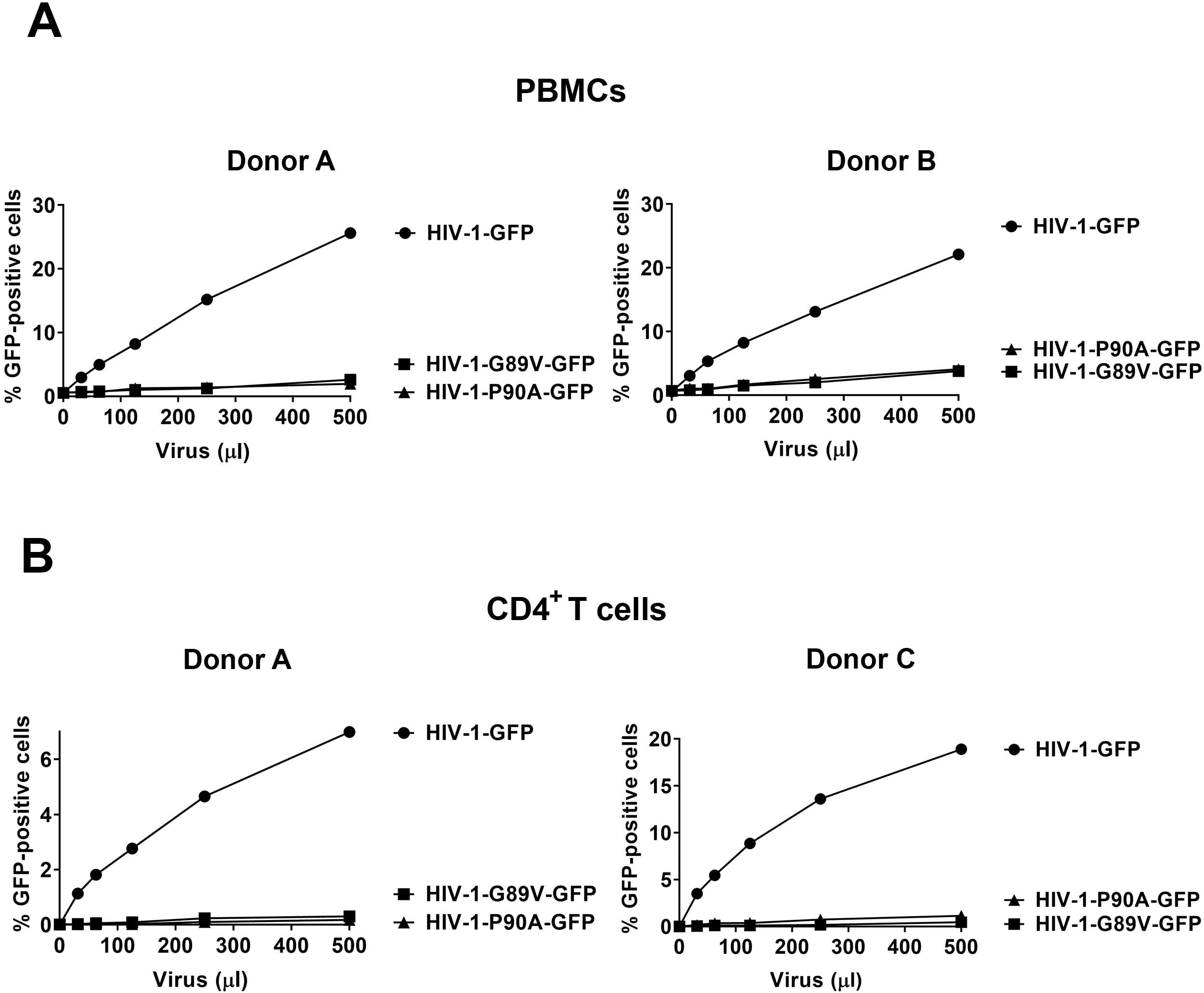
CypA-capsid Interactions are required for HIV-1 infection of human peripheral blood mononuclear cells (PBMCs) and CD4^+^ T cells. **(A)** PBMCs from two different donors were challenged with increasing amounts of HIV-1-GFP, HIV-1-G89V-GFP, or HIV-1-P90A-GFP. Infection was determined at 72 h post-challenge by measuring the percentage of GFP-positive cells. Experiments were repeated three times per donor and a representative experiment is shown. **(B)** Purified CD4^+^ T cells from two different donors were challenged with increasing amounts of HIV-1-GFP, HIV-1-G89V-GFP, or HIV-1-P90A-GFP. Infection was determined at 72 h post-challenge by measuring the percentage of GFP-positive cells. Experiments were repeated three times per donor and a representative experiment is shown.

### Disruption of HIV-1 binding to TRIM5α_hu_ does not affect viral infection in human PBMCs and CD4^+^ T cells

Mutant HIV-1 bearing the capsid change A92E lacks the ability to interact with TRIM5α_hu_. We examined the ability of HIV-1-A92E to infect PBMCs and CD4^+^ T cells to study the role of TRIM5α_hu_ during viral infection in these cells. Consistent with the results observed in Jurkat cells, HIV-1-A92E infection levels were comparable to that of wild-type HIV-1 in human PBMCs (Figure 7A). As a control, we also challenged PBMCs with increasing concentrations of HIV-1-P90A, an HIV-1 mutant that is completely inhibited in PBMCs (Figure 7A). We observed similar results when we challenged CD4^+^ T cells using HIV-1-A92E (Figure 7B). Consistent with our previous binding experiments, we found that HIV-1-A92E infection of PBMCs and CD4^+^ T cells was not affected by TRIM5α_hu_, which correlated with the inability of TRIM5α_hu_ to bind to mutant HIV-1-A92E cores.

**Figure 7.**
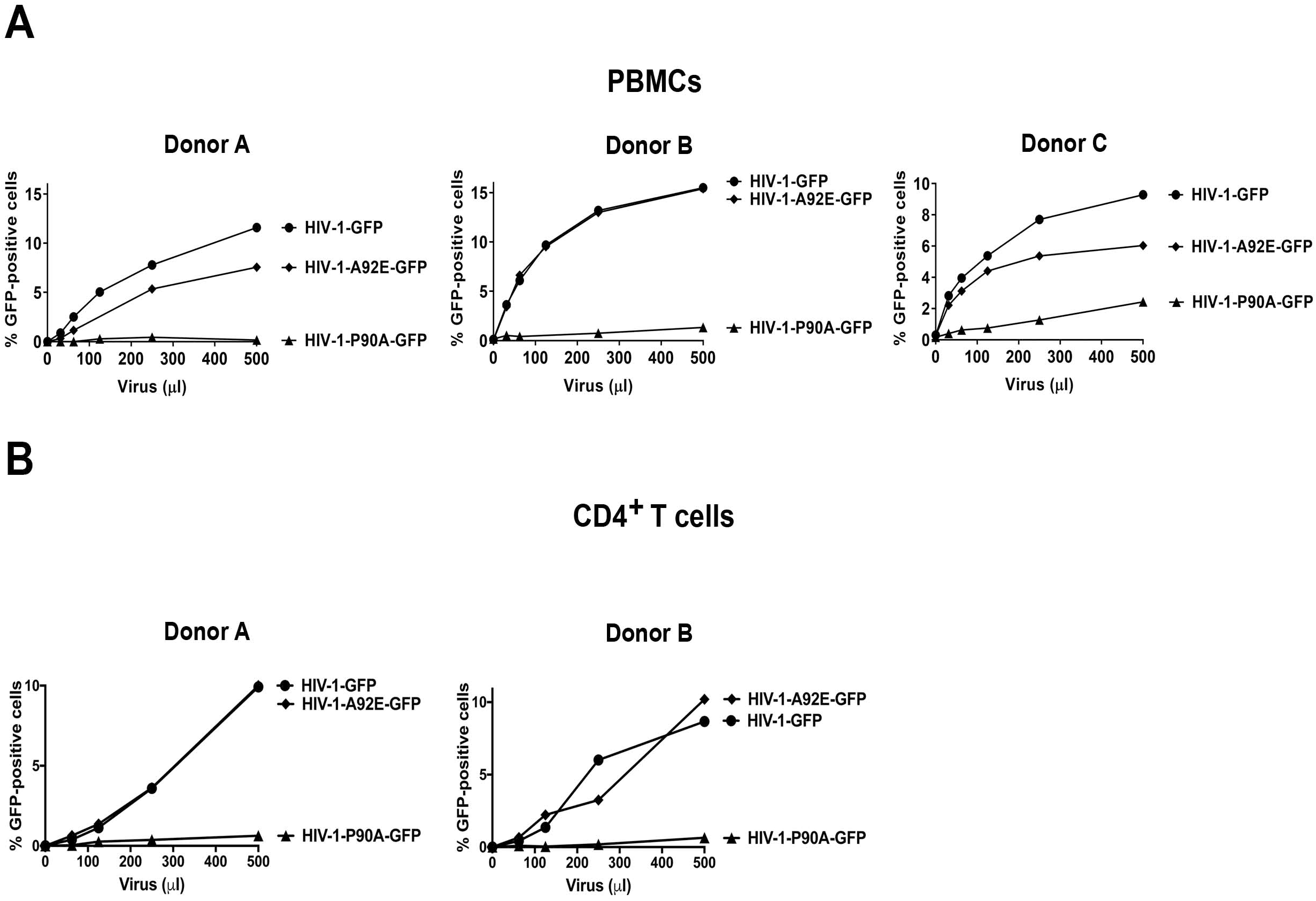
Infection of HIV-1 bearing the capsid change A92E, which does not bind TRIM5α_hu_, is not affected in human PBMCs and CD4^+^ T cells. **(A)** PBMCs from three different donors were challenged with increasing amounts of HIV-1-GFP, HIV-1-P90A-GFP, or HIV-1-P92E-GFP. Infection was determined at 72 h post-challenge by measuring the percentage of GFP-positive cells. Experiments were repeated three times per donor and a representative experiment is shown. **(B)** CD4^+^ T cells from three donors were challenged with increasing amounts of HIV-1-GFP, HIV-1-P90A-GFP, or HIV-1-P92E-GFP. Infection was determined at 72 h post-challenge by measuring the percentage of GFP-positive cells. Experiments were repeated three times per donor and a representative experiment is shown.

### Depletion of CypA expression in human CD4^+^ T cells inhibits HIV-1 infection

We have previously shown that CypA-capsid interactions are important for HIV-1 infection. To directly test the role of CypA in viral infection, we challenged CypA knockout CD4^+^ T cells with HIV-1_NL4-3_-GFP. To create CypA knockouts, CD4^+^ T cells were separately electroporated with CRISPR/Cas9 complexes containing five different guides against CypA (CypA-KO#1–5). Expression of CypA was analyzed by western blotting using anti-CypA antibodies at 3 days post-electroporation. As shown in Figure 8A, all five guides were able to eliminate the expression of CypA. The CypA knockout cells were then challenged with HIV-1_NL4-3_-GFP. Consistent with our previous results, human CD4^+^ T cells depleted for CypA expression were resistant to HIV-1 infection (the results from two donors are shown in Figure 8B). As a control, we used a CRISPR/Cas9 complex containing an RNA guide for CXCR4, which completely inhibits infection of HIV-1_NL4-3_-GFP. We also used a CRISPR/Cas9 complex containing a non-targeting RNA guide. These results demonstrated that CypA expression is essential for HIV-1 infection in human primary CD4^+^ T cells.

**Figure 8.**
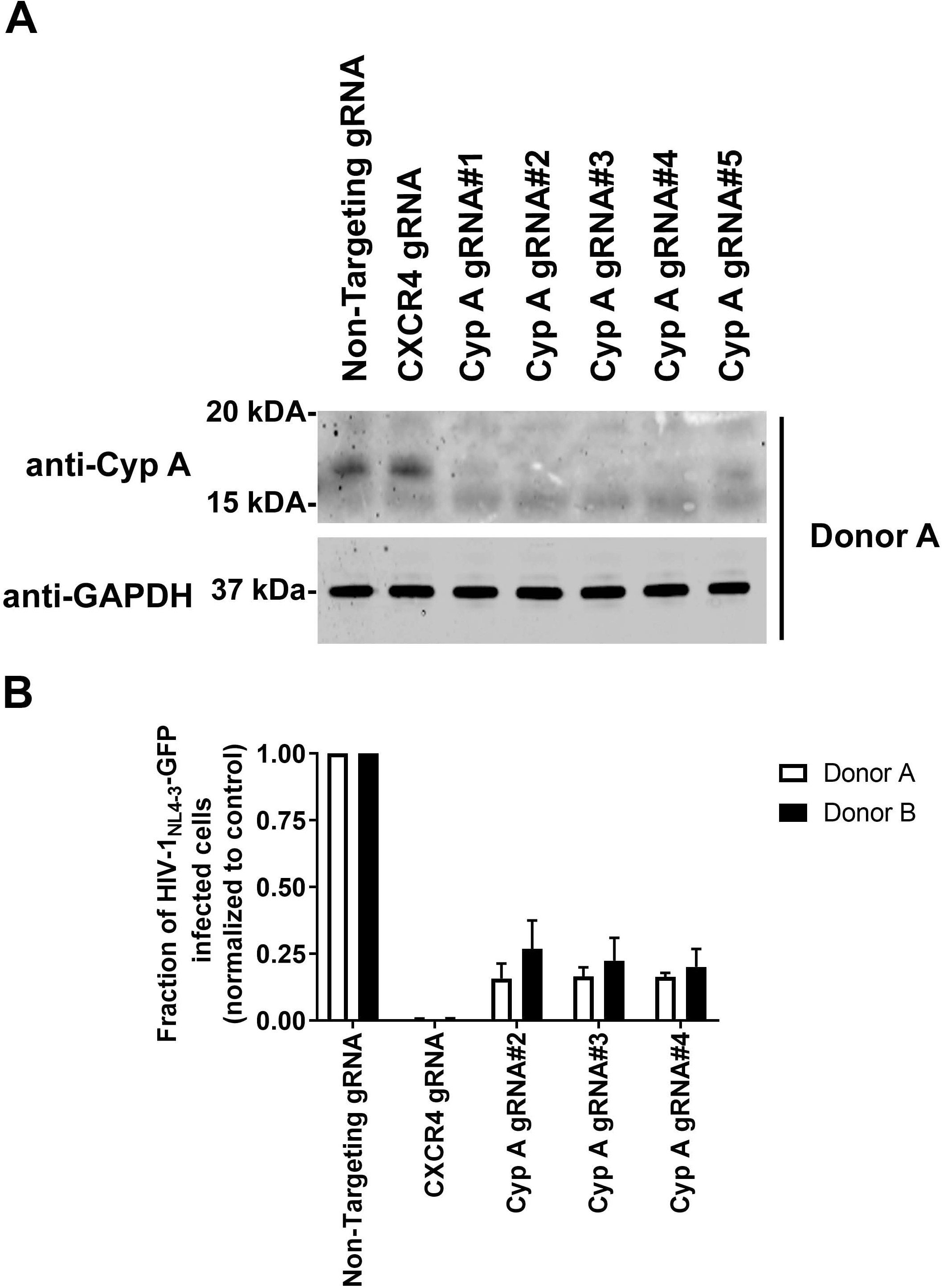
Depletion of CypA expression in human CD4^+^ T cells inhibits HIV-1 infection. **(A)** Human primary CD4^+^ T cells from two different donors were knocked-out for CypA expression using the CRISPR-Cas9 system, as described in methods. Briefly, CD4^+^ T cells were electroporated using five different guide RNAs (gRNAs) against CypA (gRNA#1-5) together with the Cas9 protein. At 72 h post-electroporation, endogenous expression of CypA in CD4^+^ T was analyzed by western blotting using antibodies against CypA. As controls, we electroporated a gRNA against CXCR4 and a non-targeting gRNA. Expression of GAPDH was used as a loading control. Similar results were obtained using two different donors and a representative blot is shown. **(B)** Human primary CD4^+^ T cells depleted for CypA expression were challenged with a replication-competent HIV-1 expressing GFP as a reporter of infection (HIV-1_NL4-3_-GFP). Infection was determined at 72 h post-infection by measuring the percentage of GFP-positive cells. Experiments were performed in triplicates, and results for two different donors with standard deviation are shown.

### CypA-capsid interactions are required for efficient HIV-1 reverse transcription in human CD4^+^ T cells

Our analysis showed that HIV-1 strains bearing capsid mutations that disrupt CypA-capsid interactions are unable to infect human CD4^+^ T cells. To identify the step in the HIV-1 life cycle at which infection is disrupted, we challenged human CD4^+^ T cells with p24-normalized levels of GFP expressed by wild-type and capsid mutant HIV-1 particles, and measured reverse transcription, nuclear import, and integration using real-time polymerase chain reaction (PCR) (Figure 9). Infection was measured by determining the percentage of GFP-positive cells at 72 h post-infection (Figure 9A). Consistent with our previous results, HIV-1-P90A and HIV-1-G89V strains showed weak infection in CD4^+^ T cells (Figure 9A). Reverse transcription was measured by determining the levels of late reverse transcripts (LRT) at 10 h post-infection (Figure 9B). Interestingly, reverse transcription was reduced in HIV-1-P90A and HIV-1-G89V, suggesting that CypA-capsid interactions were important for HIV-1 reverse transcription in CD4^+^ T cells. As a control, we used Nevirapine (Nev), which inhibits reverse transcription (Figure 9B). For nuclear import, we measured the levels of 2-long terminal repeat (2-LTR) circles at 24 h post-infection (Figure 9C). Consistent with a reverse transcription block, our results showed that both viruses were blocked for nuclear import. Viral integration was determined using Alu-PCR at 48 h post-infection (Figure 9D). As a control, the integration inhibitor Raltegravir (Ral) was also used. As expected, the integration of HIV-1-P90A and HIV-1-G89V was blocked. Overall, these results suggested that HIV-1 requires interaction with CypA in order to undergo efficient reverse transcription in human CD4^+^ T cells. These results also suggested that in the absence of CypA, HIV-1 particles are exposed to TRIM5α_hu_, which inhibits reverse transcription.

**Figure 9.**
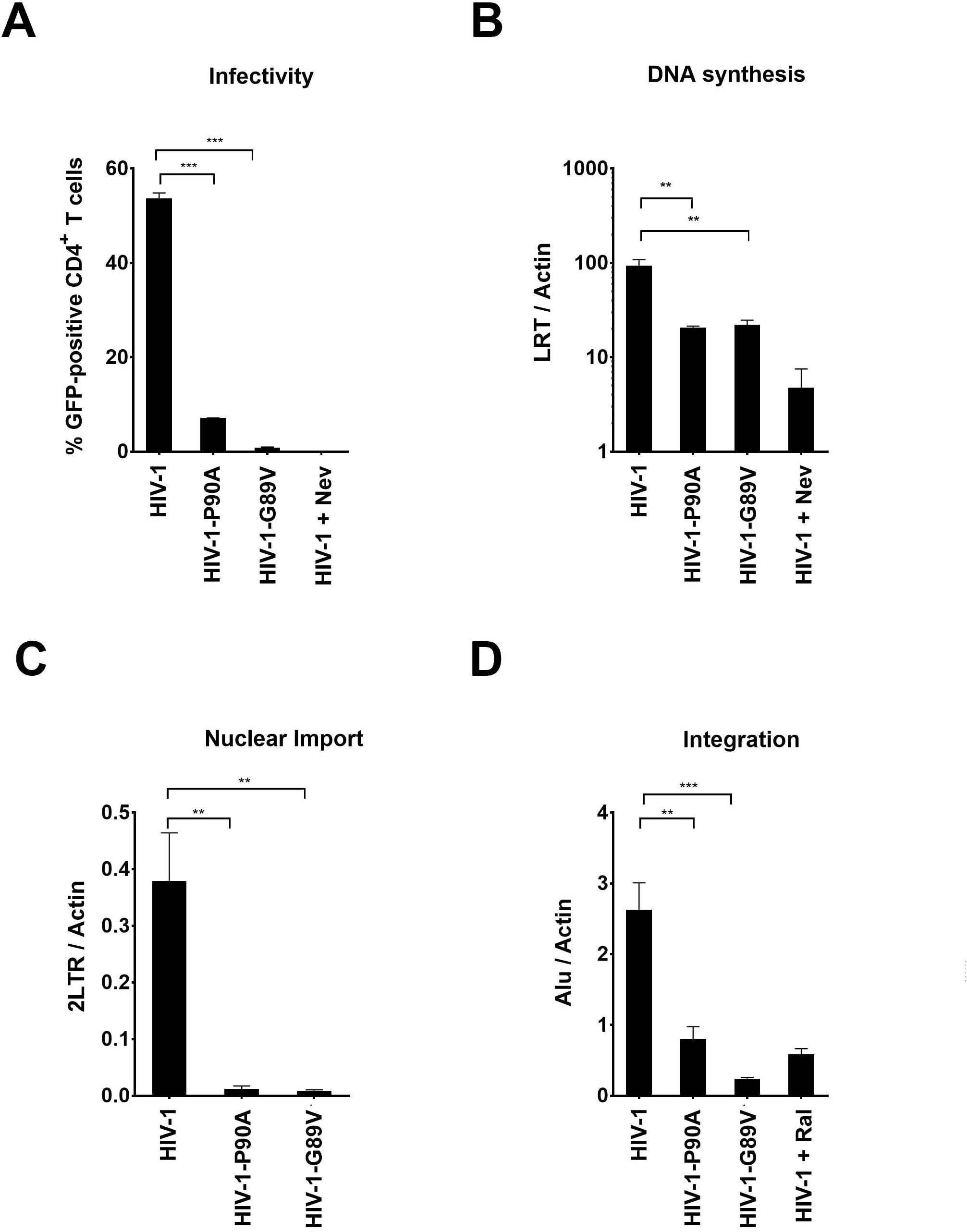
CypA-capsid interactions are required for HIV-1 reverse transcription in human CD4^+^ T cells. Human primary CD4^+^ T cells were challenged with DNase-pretreated HIV-1-GFP, HIV-1-P90A-GFP, or HIV-1-G89V-GFP at a multiplicity of infection (MOI) < 1.0. Infection was determined by measuring the percentage of GFP-positive cells using flow cytometry at 72 h post-challenge **(A)**. In parallel, total DNA was extracted from similar infections at 12, 36, and 72 h post-infection. The DNA samples collected at 12 h post-infection were used to determine the levels of late reverse transcripts (LRTs) using real-time PCR **(B)**, as described in methods. As a control, we inhibited reverse transcription by using 10 µm of Nevirapine (Nev). HIV-1 2-LTR circles, which is an indirect measure of nuclear import, were quantified using real-time PCR in DNA samples collected 36 h post-infection **(C)**, as described in methods. Integrated proviruses were measured by determining the amount of Alu-PCR (Alu) products using real-time PCR in DNA samples collected at 72 h post-infection **(D)**, as described in methods. As a control, we inhibited proviral integration by using 10 µM Raltegravir (Ral). LRT, 2-LTR, and Alu products were normalized to actin. Results were analyzed using the unpaired t-test. Differences were considered statistically significant at P < 0.01 (**), P < 0.001 (***), P < 0.0001 (****).

## DISCUSSION

A few years after the discovery of HIV-1 capsid interaction with CypA (Luban et al., 1993), it was established that CsA treatment disrupted this interaction, which then affected HIV-1 infectivity. It was soon realized that CsA exerted varying effects on HIV-1 infection depending upon the human cell lines used. Although CypA-capsid interactions have been studied for over 25 years, the mechanism by which CsA affects HIV-1 infection remains elusive. Here, we explored the role of CypA-capsid interactions in human lymphocytes.

Several recent studies have suggested that TRIM5α_hu_ may function as an HIV-1 restriction factor (Jimenez-Guardeno et al., 2019; OhAinle et al., 2018). This led us to hypothesize that TRIM5α_hu_ binds to the HIV-1 core, which was proven by our results. Remarkably, we observed that TRIM5α_hu_ showed stronger binding to the HIV-1 core than TRIM5α_rh_. Because TRIM5α_hu_ and CypA both bind to the HIV-1 core, it is possible that these two factors compete for the HIV-1 core during infection⎯CypA may modulate the binding of TRIM5α_hu_ to the HIV-1 core or vice versa. We showed that the depletion of TRIM5α_hu_ expression rescued the HIV-1 infectivity phenotype in Jurkat cells caused by the disruption of CypA-capsid interactions. In cellular extracts, we also observed that CsA treatment increased binding of TRIM5α_hu_ to the HIV-1 core. These results were consistent with our hypothesis that CypA modulates the binding of TRIM5α_hu_ to the HIV-1 core. The results of the binding experiments suggested that CypA sterically hinders the binding of TRIM5α_hu_ to the HIV-1 core, which could explain the observed infectivity phenotype when CypA-capsid interactions were disrupted in lymphocytes. Conversely, we also demonstrated that increasing the amounts of CypA prevented the binding of TRIM5α_hu_ to the HIV-1 core. Based on these results, we propose that the prevention of CypA binding allows TRIM5α_hu_ to bind to the HIV-1 core, consequently inhibiting infection.

Our results indicated that the interaction of TRIM5α_hu_ with the HIV-1 core leads to restriction of infection. However, several previous studies have shown that overexpression of TRIM5α_hu_ has little or no effect on HIV-1 infection (Li et al., 2006; Pham et al., 2010; Yap et al., 2005). One possibility is that restriction of HIV-1 infection by TRIM5α_hu_ requires a specific cellular cofactor expressed only in Jurkat cells and human primary lymphocytes. This cofactor may have been absent or present only in insufficient amounts in other human cell lines that were used for TRIM5α_hu_ overexpression experiments. The search for this type of cofactors in the future is warranted. Alternatively, TRIM5α_hu_ might be recruited to the HIV-1 core by other TRIM proteins through the formation of higher-order complexes (Diaz-Griffero et al., 2009); TRIM5α dimers are known to associate with other dimers and form higher-order self-association complexes. For example, TRIM5α_rh_ forms higher-order complexes with human TRIM34 and TRIM6 (Li et al., 2011). Alternatively, TRIM5α_hu_ might recruit other human TRIM orthologs that are important for restriction. Future studies will explore the existence of other human TRIM orthologs that may form higher-order complexes with TRIM5α_hu_.

Mutations in the CypA-binding loop of the capsid, such as P90A and G89V, cause an infectivity defect similar to that caused by CsA treatment during wild-type HIV-1 infection of Jurkat cells; however, infection of HIV-1 viruses bearing the capsid change A92E are insensitive to CsA treatment in Jurkat cells (Hatziioannou et al., 2005; Sokolskaja et al., 2004). Here, we showed that this insensitivity was due to the lack of TRIM5α_hu_ binding to HIV-1-A92E mutant cores. Normal viral infectivity correlated with the inability of TRIM5α_hu_ to bind to HIV-1-A92E cores. Additionally, the HIV-1-A92E mutant core also did not interact with CypA, indicating that the A92E mutation abolishes interaction of the HIV-1 core with both TRIM5α_hu_ and CypA. Based on these results, it is possible that TRIM5α_hu_ binds to the CypA-binding loop or in its proximity.

The disruption of CypA-capsid interactions had a stronger effect on viral infectivity in human primary cells compared with Jurkat cells. Mutant HIV-1 (P90A or G89V) infection of PBMCs or CD4^+^ T cells was 10–20 fold weaker than that of wild-type HIV-1. Similarly, KD of CypA expression in human primary CD4^+^ T cells potently blocked HIV-1 infection. In contrast, HIV-1-A92E infection of primary lymphocytes was not affected when compared to that of wild-type HIV-1. Taken together, these results suggested that TRIM5α_hu_ blocks HIV-1 infection of human primary lymphocytes when CypA-capsid interactions are disrupted. Consistent with this idea, we observed that the inhibition of infection caused by the disruption of CypA-capsid interactions in CD4^+^ T cells occurred prior to HIV-1 reverse transcription.

Our work has shown that CypA protects the core from TRIM5α_hu._ Intriguingly, new- and old-world monkeys express a protein in which the TRIM domain of TRIM5α and CypA form a fusion protein (Brennan et al., 2008; Newman et al., 2008; Sayah et al., 2004). Based on our results, it is possible that evolution may have taken advantage of a positive factor, in this case CypA, and fused to the TRIM domain to develop a restriction factor.

Overall, this work showed the mechanism by which disruption of CypA-capsid interactions affect HIV-1 infection of human primary lymphocytes. Future experiments will explore whether a similar mechanism operates in macrophages or dendritic cells.

## ACKNOWLEDGEMENTS

We thank the NIH AIDS repository for important reagents. We are very grateful to Kyusik Kim and Jeremy Luban for reagents and helpful discussions. This work was supported by an R01 grant from the NIH AI087390, to F.D.-G.

## MATERIALS AND METHODS

### Cell lines and drugs

Human Jurkat lymphocytes cells obtained from the American type culture collection (ATCC) were grown at 37°C in Roswell Park Memorial Institute (RPMI) medium supplemented with 10% fetal bovine serum (FBS) and 1% penicillin-streptomycin. Human 293T (ATCC) cells were grown at 37°C in Dulbecco’s modified eagle medium (DMEM) supplemented with 10% FBS and 1% penicillin-streptomycin. Cyclosporin A (CsA) was dissolved in dimethyl sulfoxide (DMSO) to create a stock of 10 mM.

### Infection using HIV-1-GFP reporter viruses

Recombinant human immunodeficiency virus-1 (HIV-1) strains such as HIV-1-G89V, HIV-1-P90A, and HIV-1-A92E expressing GFP were prepared as previously described (Diaz-Griffero et al., 2008). Viral challenges were performed in 24-well plates by infecting 50,000 cells (Jurkat, PBMCs, or CD4^+^ T cells) per well. Infection was determined by measuring the percentage of GFP-positive cells using flow cytometry (Becton Dickinson).

### Capsid expression and purification

pET-11a vectors were used to express the HIV-1 capsid protein containing the A14C and E45C mutations. Point mutations P90A and A92E were introduced using the QuikChange II site-directed mutagenesis kit (Stratagene) according to the manufacturer’s instructions. All proteins were expressed in *Escherichia coli* one-shoot BL21star (DE3) cells (Invitrogen), as previously described (Selyutina et al., 2018). Briefly, cells were inoculated in Luria-Bertani (LB) medium and cultured at 30°C until mid-log phase (*A*_600_, 0.6–0.8). Protein expression was induced with 1 mM isopropyl-β-d-thiogalactopyranoside (IPTG) overnight at 18°C. Cells were harvested by centrifugation at 5,000 × g for 10 min at 4°C, and pellets were stored at −80°C until purification. Purification of capsids was carried out as follows: pellets from 2 L of Luria containing bacteria were lysed by sonication (Qsonica microtip: 4420; A = 45; 2 min; 2 sec on; 2 sec off for 12 cycles), in 40 ml of lysis buffer (50 mM Tris pH = 8, 50 mM NaCl, 100 mM β-mercaptoethanol, and Complete EDTA-free protease inhibitor tablets). Cell debris was removed by centrifugation at 40,000 × g for 20 min at 4°C. Proteins from the supernatant were precipitated by incubation with one-third the volume of saturated ammonium sulfate containing 100 mM β-mercaptoethanol for 20 min at 4°C, and centrifugation at 8,000 × g for 20 min at 4°C. Precipitated proteins were resuspended in 30 ml of buffer A (25 mM 2-(N-morpholino) ethanesulfonic acid (MES) pH 6.5 and 100 mM β-mercaptoethanol) and sonicated 2–3 times (Qsonica microtip: 4420; A = 45; 2 min; 1 sec on; 2 sec off). The protein sample was dialyzed three times in buffer A (2 h, overnight, 2 h), sonicated, diluted in 500 ml of buffer A, and was chromatographed sequentially on a 5-ml HiTrap Q HP column and on a 5-ml HiTrap SP FF column (GE Healthcare), which were both pre-equilibrated with buffer A. The capsid protein was eluted from the HiTrap SP FF column using a linear gradient of concentrations ranging from 0–2 M NaCl. The eluted fraction that had highest protein levels was selected based on absorbance at 280 nm. Pooled fractions were dialyzed three times (2 h, overnight, 2 h) in storage buffer (25 mM MES, 2 M NaCl, 20 mM β-mercaptoethanol). The sample was reduced to a concentration of 20 mg/ml using Centricon and stored at −80°C.

### Assembly of stabilized HIV-1 capsid tubes

One milliliter of monomeric capsid (5 mg/ml) was dialyzed in SnakeSkin dialysis tubing 10K MWCO (Thermo Scientific) using a buffer that is high in salt and contains a reducing agent (buffer 1: 50 mM Tris, pH 8, 1 M NaCl, 100 mM β-mercaptoethanol) at 4°C for 8 h. Subsequently the protein was dialyzed using buffer 1 without the reducing agent β-mercaptoethanol (buffer 2: 50 mM Tris, pH 8, 1 M NaCl) at 4°C for 8 h. The absence of β-mercaptoethanol in the second dialysis allows the formation of disulfide bonds between Cysteine 14 and 45 inter-capsid monomers in the hexamer. Finally, the protein was dialyzed using buffer 3 (20 mM Tris, pH 8,0, 40 mM NaCl) at 4°C for 8 h. Assembled complexes were kept at 4°C for up to 1 month.

### Capsid binding assay protocol

Human HEK293T cells were transfected for 24 h with a plasmid expressing the protein of interest (TRIM5α_hu_,TRIM5α_rh_, or CypA). The cell medium was completely removed and cells were scraped off the plate and lysed in 300 μl of capsid binding buffer (CBB: 10 mM Tris, pH 8,0, 1,5 mM MgCl_2_, 10 mM KCl). Cells were rotated at 4°C for 15 min and then centrifuged to remove cellular debris (21,000 *×* g, 15 min, 4°C). Cell lysates were incubated with stabilized HIV-1 capsid tubes for 1 h at 25°C. Subsequently, the stabilized HIV-1 capsid tubes were centrifuged at 21,000 × g for 2 min. Pellets were washed 2-3 times by resuspension and centrifugation in CBB or phosphate-buffered saline (PBS). Pellets were resuspended in Laemmli buffer 1× and analyzed by western blot using anti-p24 and the indicated antibodies.

### Preparation of PBMCs and CD4^+^ T cells

PBMCs from healthy donor whole blood were isolated by density gradient centrifugation using Ficoll-Paque Plus (GE Health Care, #17-1440-02). 40 ml of whole blood was centrifuged at 300 × g for 10 min, and the plasma layer was removed and replaced with Hank’s Balanced Salt solution (HBSS; Sigma Aldrich, #H6648). The whole blood was then diluted to a ratio of 1:2 with HBSS with 20 ml of the diluted blood layered on top of 20 ml Ficoll-Paque Plus and centrifuged at 300 × g for 30 min. The resulting whitish buffy coat layer was collected, washed twice with HBSS, and resuspended in 10 RPMI medium containing 10% (vol/vol) FBS and 1% (vol/vol) penicillin-streptomycin, and activated with IL-2 (100 U/ml) (Human IL-2; Cell Signaling Technology, #8907SF) and phytohemagglutinin (1 µg/ml) for 3 days. CD4^+^ T cells were obtained via negative selection from PBMCs using a human CD4^+^ T-cell isolation kit (MACS Miltenyi Biotec, #130-096-533). Per 1 × 10^7^ total cells, PBMCs were resuspended in 40 µl CD4^+^ T-cell isolation buffer (PBS, pH 7.2, 0.5% bovine serum albumin, and 2 mM EDTA). Ten microliters of CD4^+^ T-cell biotin-antibody cocktail was then added and incubated at 4°C for 5 min. Thirty microliters of CD4^+^ T-cell isolation buffer and 20 µl of CD4^+^ T-cell microbead cocktail were then added and further incubated for 10 min at 4°C. Depending on the amount of PBMCs isolated, LS Column (MACS Miltenyi Biotec, #130-042-401) or MS Column (MACS Miltenyi Biotec, #130-042-201) attached to a MACS Separator was pre-washed with 3 ml or 6 ml of ice-cold CD4^+^ T-cell isolation buffer, respectively. PBMC suspension was added to the column and the flow-through was collected in a 15-ml tube. The LS or MS Columns were then washed with 3 ml or 6 ml of ice-cold CD4^+^ T-cell isolation buffer, respectively, and the flow-through was collected. The newly isolated CD4^+^ T cells were then centrifuged at 800 *x g* for 5 min and resuspended in RPMI medium supplemented with IL-2 (100 U/ml).

### Knockdown of TRIM5α_hu_ expression in Jurkat cells

Lentiviral particles were produced in human 293T cells by co-transfection of 1 µg TAT, 1 µg REV, 1.5 µg VSVG, 1 µg HDM(codon optimized gag-pol), and 7.5 µg of the lentiviral vector pLKO1, or 7.5 µg the pLKO1 vector containing an shRNA that targets TRIM5α_hu_ (pLKO1-shRNA-TRIM5α_hu_). Viruses were harvested at 48 h post-transfection and used to transduce Jurkat cells for 5 days at a 1:1 virus to complete media ratio. Transduced cells were selected in 0.15 µg/ml of puromycin. Cells that stably contained pLKO1 or pLKO1-shRNA-TRIM5α_hu_ were maintained in puromycin. Expression of TRIM5α_hu_ was evaluated by challenging the cells with N-tropic murine leukemia virus (N-MLV) and B-tropic murine leukemia virus (B-MLV).

### CRISPR-Cas9 RNP Production, CD4+ T Cell Isolation, and Electroporation

Detailed protocols for RNP production and primary CD4+ T cell editing have been previously published (Hultquist et al., 2016). Briefly, lyophilized crRNA and tracrRNA (Dharmacon) were suspended at a concentration of 160 µM in 10 mM Tris-HCL (7.4 pH) with 150 mM KCl. 5µL of 160µM crRNA was mixed with 5µL of 160µM tracrRNA and incubated for 30 min at 37°C. The gRNA:tracrRNA complexes were then mixed gently with 10µL of 40µM Cas9 (UC-Berkeley Macrolab) to form Cas9 ribonucleoproteins (RNPs). Five 3.5µL aliquots were frozen in Lo-Bind 96-well V-bottom plates (E&K Scientific) at −80°C until use. All crRNA guide sequences used in this study were derived from the Dharmacon pre-designed Edit-R library for gene knock-out, including the non-targeting (U-007502, U-007501) and CYPA-targeting guides (CM-004979-01 through CM-004979-05).

Primary human CD4+ T cells from healthy donors were isolated from donated leukoreduction chambers after Trima Apheresis (Blood Centers of the Pacific). Peripheral blood mononuclear cells (PBMCs) were isolated by Ficoll centrifugation using SepMate tubes (STEMCELL, per manufacturer’s instructions). Bulk CD4+ T cells were subsequently isolated from PBMCs by magnetic negative selection using an EasySep Human CD4+ T Cell Isolation Kit (STEMCELL, per manufacturer’s instructions). Isolated CD4+ T cells were suspended in complete Roswell Park Memorial Institute (RPMI), consisting of RPMI-1640 (Sigma) supplemented with 5mM 4-(2-hydroxyethyl)-1-piperazineethanesulfonic acid (HEPES, Corning), 2mM Glutamine (UCSF Cell Culture Facility), 50μg/mL penicillin/streptomycin (P/S, Corning), 5mM sodium pyruvate (Corning), and 10% fetal bovine serum (FBS, Gibco). Media was supplemented with 20 IU/mL IL-2 (Miltenyi) immediately before use. For activation, bulk CD4+ T cells were immediately plated on anti-CD3 coated plates [coated for 2 hours at 37°C with 20µg/mL anti-CD3 (UCHT1, Tonbo Biosciences)] in the presence of 5µg/mL soluble anti-CD28 (CD28.2, Tonbo Biosciences). Cells were stimulated for 72 hours at 37°C / 5% CO2 prior to electroporation.

Each electroporation reaction consisted of 4×10^5 T cells, 3.5 µL RNP, and 20 µL electroporation buffer. After three days of stimulation, cells were suspended and counted. RNPs were thawed and allowed to come to room-temperature. Immediately prior to electroporation, cells were centrifuged at 400xg for 5 minutes, supernatant was removed by aspiration, and the pellet was resuspended in 20 µL of room-temperature P3 electroporation buffer (Lonza) per reaction. 20 µL of cell suspension was then gently mixed with each RNP and aliquoted into a 96-well electroporation cuvette for nucleofection with the 4D 96-well shuttle unit (Lonza) using pulse code EH-115. Immediately after electroporation, 80 µL of pre-warmed media without IL-2 was added to each well and cells were allowed to rest for at least one hour in a 37°C cell culture incubator. Cells were subsequently moved to 96-well flat-bottomed culture plates pre-filled with 100 µL warm complete media with IL-2 at 40 IU/mL (for a final concentration of 20 IU/mL) and anti-CD3/anti-CD2/anti-CD28 beads (T cell Activation and Stimulation Kit, Miltenyi) at a 1:1 bead:cell ratio. Cells were cultured at 37°C / 5% CO_2_ in a dark, humidified cell culture incubator for 4 days to allow for gene knock-out and protein clearance, with additional media added on day 2. To check CYPA knock-out efficiency, 50 µL of mixed culture was removed to a centrifuge tube. Cells were pelleted, supernatant was removed, and pellets were resuspended in 100 µL 2.5x Laemmli Sample Buffer. Protein lysates were heated to 98°C for 20 min before storage at −20°C until Western blotting (below).

### Detection of Late Reverse Transcripts, 2-LTR circles, and integration in CD4^+^ T cells

Total DNA was extracted from 1 × 10^6^ infected cells. Samples were processed at 12, 36, and 66 h post-infection using QIAmp DNA micro kit (QIAGEN). Viral DNA forms of wild-type and mutant HIV-1 were amplified using real-time PCR (Butler et al., 2001; Valle-Casuso et al., 2012). β-actin amplification was used for normalization. LRT, which refers to the whole synthesis of DNA by HIV-1, were measured at 12 h post-infection. 2-LTR circles were measured at 36 h post-infection to assess nuclear import. Provirus integration in cellular genome was measured at 66 h post-infection using Alu-PCR. Reactions were performed in 1X FastStart Universal Probe Master Mix (Rox) 2× (Roche) in 20 µl volume. The PCR reaction consisted of the following steps: initial annealing (50°C 2 min), denaturation step (95°C 15 min), 40 cycles of amplification (95°C 15 sec, 58°C 30 sec, 72°C 30 sec). Primer or probe sequences are as follows: LRT MH531: 5’-TGTGTGCCCGTCTGTTGTGT-3’; MH532: 5’-GAGTCCTGCGTCGAGAGAGC-3’; LRT-Probe: 5′-(FAM)-CAGTGGCGCCCGAACAGGGA-(TAMRA)-3’. 2-LTR circles: MH535: 5’-AACTAGGGAACCCACTGCTTAAG-3’; MH536: 5′-TCCACAGATCAAGGATATCTTGTC-3′; 2-LTR probe: MH603: 5’-(FAM)-ACACTACTTGAAGCACTCAAG-GCAAGCTTT-(TAMRA)-3’. Alu-PCR 1^st^ round: Alu 1: 5’-TCCCAGCTACTCGGGAGGCTGAGG-3’; Alu 2: 5’-GCCTCCCAAAGTGCTGGGATTACAG-3’; Lambda U3: 5′-ATGCCACGTAAGCGAAACTTTCCGCTGGGGACTTTCCAGGG-3′. Alu quantitative PCR 2^nd^ round: Lambda: (5’-ATGCCACGTAAGCGAAACT-3’); U5: (5’-CTGACTAAAAGGGTCTGAGG-3’; Probe: 5’-(FAM)-TTAAGCCTCAATAAAGCTTGCCTTGAGTGC-(TAMRA). β-actin: FX: 5’-AACACCCCAGCCATGTACGT-3’, RX: 5-CGGTGAGGATCTTCATGAGGTAGT, Actin-Probe: (6FAM)-CCAGCCAGGTCCAGACGCAGGA-(BBQ).

### Western blot analysis

Cells were lysed in lysis buffer (50 mM Tris [pH 8.0], 280 mM NaCl, 0.5% IGEPAL 40, 10% glycerol, 5 mM MgCl_2_). Proteins were detected by western blot using anti-FLAG (1:1,000 dilution; Sigma), anti-hemagglutinin (HA) (1:1,000 dilution; Sigma), anti-CypA (1:5,000 dilution; catalog number ab58144; Abcam), or anti-glyceraldehyde-3-phosphate dehydrogenase (GAPDH) (1:5,000; Ambion), or anti-p24 (1:1,000, catalog number 183-H12-5C; NIH) antibodies. Secondary antibodies against rabbit and mouse IgG conjugated to IRDye 680LT or IRDye 800CW were obtained from Li-Cor (1:10,000 dilution). Bands were detected by scanning blots using the Li-Cor Odyssey imaging system in the 700-nm or 800-nm channels.

---

**Figure S1.**
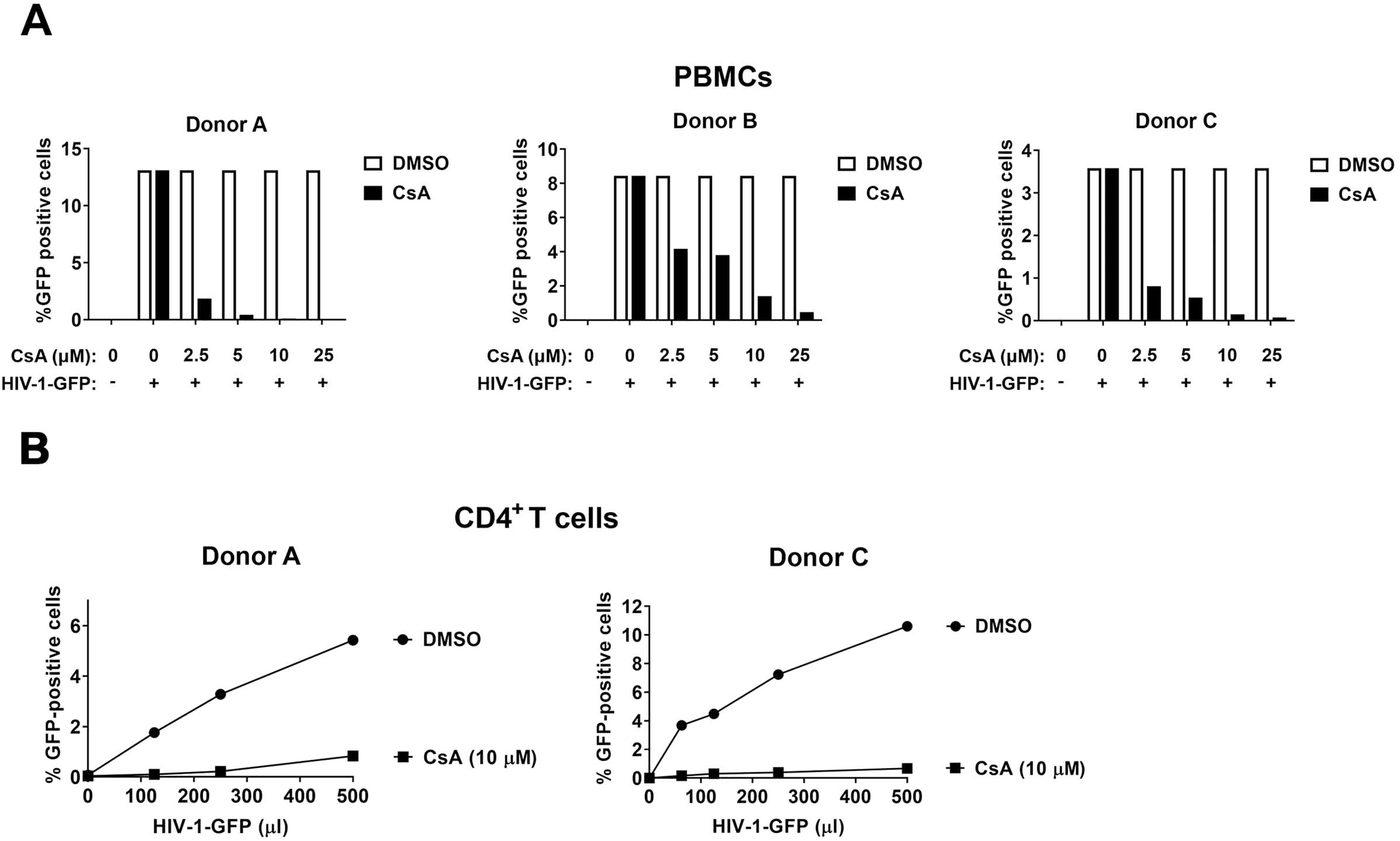
Inhibition of CypA-capsid interactions with CsA restricts HIV-1 infection in PBMCs and CD4^+^ T cells. **(A)** PBMCs from three different donors were challenged with HIV-1-GFP in the presence of increasing concentrations of CsA. DMSO was used as a control. Infection was determined at 72 h post-challenge by measuring the percentage of GFP-positive cells. **(B)** CD4^+^ T cells from two different donors were challenged with HIV-1-GFP in the presence of 10 µM CsA. DMSO was used as a control. Infection was determined at 72 h post-challenge by measuring the percentage of GFP-positive cells.

